# *STM* is required for fate establishment of productive shoot progenitor cells in *Arabidopsis* tissue culture

**DOI:** 10.1101/2025.04.14.648630

**Authors:** Ning Zhai, Dixiang Xie, Lin Xu

## Abstract

De novo shoot regeneration via the two-step tissue culture method is widely used for vegetative reproduction of plants and for gene transformation and editing. A key step in tissue culture is the fate transition from shoot progenitor cells (SPCs) to productive SPCs (prSPCs), which are capable of differentiating into a shoot. In this study, we identified the class I KNOTTED1-like homeobox (KNOX) transcription factor gene *SHOOT-MERISTEMLESS* (*STM*) as the key controller of, and marker gene for, the fate transition from SPCs into prSPCs, indicating that *STM* might serve as a candidate of molecular tools to improve de novo shoot regeneration in the agricultural applications.

---

*De novo* shoot regeneration *via* the two-step tissue culture method is widely used for vegetative reproduction of plants and for gene transformation and editing. A key step in tissue culture is the fate transition from shoot progenitor cells (SPCs) to productive SPCs (prSPCs), which are capable of differentiating into a shoot. In this study, we identified the class I KNOTTED1-like homeobox (KNOX) transcription factor gene *SHOOT-MERISTEMLESS* (*STM*) as the key controller of, and marker gene for, the fate transition from SPCs into prSPCs, indicating that *STM* might serve as a candidate of molecular tools to improve *de novo* shoot regeneration in the agricultural applications.

Taking an *Arabidopsis thaliana* hypocotyl explant as an example in the two-step tissue culture (Fig. 1A), in the first step (i.e., the pluripotency acquisition step), the detached hypocotyl is cultured on auxin-rich callus-inducing medium (CIM) to produce callus. Callus is initiated from the adult vascular stem cells (e.g., procambium and xylem-pole pericycle cells) and has tissue identities similar to the root apical meristem, which is the cellular basis of the pluripotency of callus for further organogenesis (Sugimoto et al., 2010; Zhai and Xu, 2021). In the second step (i.e., the organogenesis step), callus is transferred to cytokinin-rich shoot-inducing medium (SIM) to induce shoot organogenesis. In this step, the key cell fate transition is the formation of SPCs expressing the marker gene *WUSCHEL* (*WUS*) (Zhang et al., 2017). The SPCs are located in the middle cell layer of callus and are activated by the cytokinin signaling pathway (Meng et al., 2017; Zhang et al., 2017; Zhai and Xu, 2021). Then, the cell fate of SPCs can enter two different lineages. In one lineage, a group of SPCs (but not a single SPC) changes fate to form prSPCs, which are capable of further differentiation into the pro-shoot apical meristem (proSAM) and finally the shoot apical meristem (SAM) (Varapparambath et al., 2022; Wang et al., 2025). The cell wall remodeling-induced mechanical conflict surrounding SPCs is essential for their transition into prSPCs (Varapparambath et al., 2022). In the other lineage, some SPCs lose the ability to further differentiate into the proSAM, instead becoming pseudo SPCs (psSPCs) (Varapparambath et al., 2022). However, the molecular mechanism that determines whether SPCs transition into prSPCs or psSPCs remains largely unclear.

**Figure 1.**
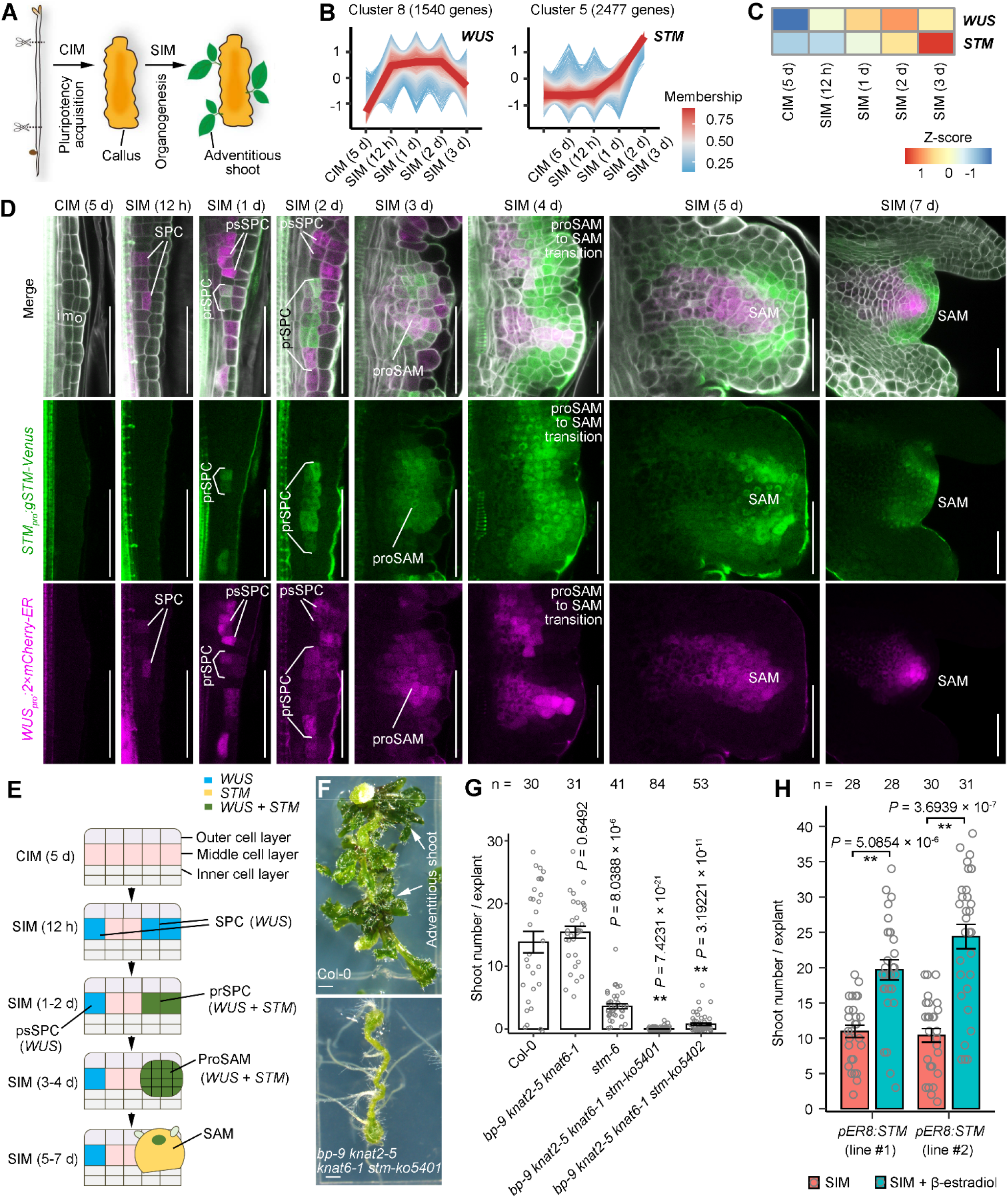
*STM* in prSPCs. **(A)** Schematic of two-step tissue culture using *Arabidopsis* hypocotyl explants. **(B)** Expression patterns of genes in clusters 8 and 6 during tissue culture, as determined from RNA-Seq data. See Figure S1 and Table S1 for full list of gene clusters. **(C)** Heat map showing transcript profiles of *WUS* and *STM* during tissue culture. **(D)** Confocal analysis of *WUS*_*pro*_*:2×mCherry-ER* and *STM*_*pro*_*:gSTM-Venus* marker lines during tissue culture. o, outer cell layer; m, middle cell layer; i, inner cell layer. **(E)** Schematic of fate transition during shoot organogenesis from callus on SIM. **(F, G)** Phenotype (F) and statistical (G) analysis of *de novo* shoot regeneration from hypocotyl explants of Col-0, *bp-9 knat2-5 knat6-1* triple mutant, *stm-6* single mutant, and *bp-9 knat2-5 knat6-1 stm-ko5401* and *bp-9 knat2-5 knat6-1 stm-ko5402* quadruple mutants. **(H)** β-estradiol-induced overexpression of *STM* (*pER8:STM*) promotes shoot regeneration from callus. The hypocotyls of *pER8:STM* were cultured on CIM for 5 days and then moved to SIM without (control) or with 5 μm β-estradiol. \*\**P* < 0.01 (Mann-Whitney U test) compared with Col-0 (G) or with control (H). Error bars show SEM (G, H). Individual values (gray dots) and means (bars) are shown in (G, H). Scale bars, 50 μm in (D) and 1 mm in (F).

To study cell fate transition during tissue culture, we cultured wild-type *Arabidopsis* Columbia-0 (Col-0) hypocotyl explants on CIM for 5 days to produce calli, and then moved the calli to SIM to induce shoot organogenesis. Calli on CIM and SIM were collected at various time points and their transcriptomes were analyzed by RNA sequencing (RNA-seq). We identified nine clusters of genes with different expression patterns in tissue culture at 5 days on CIM and at 12 hours, 1 day, 2 days, and 3 days on SIM (Supplementary Fig. S1; Supplementary Table S1). Among those clusters, cluster 8 contained genes showing low transcript levels on CIM and rapid upregulation on SIM from 12 hours onwards; this cluster contained *WUS* (Fig. 1B and 1C). *STM*, a key gene involved in SAM development (Long et al., 1996) and shoot regeneration (Endrizzi et al., 1996; Daimon et al., 2003; Zhang et al., 2017), was in cluster 6, whose members showed low transcript levels on CIM at 5 days and on SIM at 12 hours, followed by upregulation on SIM from 1 day onwards (Fig. 1B and 1C). Therefore, the upregulation of *STM* was slower than that of *WUS* in Col-0 callus cultured on SIM.

Confocal analyses of the F_3_ generation of the cross between the *WUS*_*pro*_*:2×mCherry-ER* and *STM*_*pro*_*:gSTM-Venus* marker lines showed that the expression of *WUS* and *STM* was not active at 5 days on CIM (Fig. 1D and 1E). *WUS* expression was detected at 12 hours on SIM in SPCs in the middle cell layer of callus, but *STM* expression was not detected at this stage (Fig. 1D and 1E). At 1 to 2 days on SIM, a group of SPCs started to change fate to become prSPCs, with co-expression of *STM* and *WUS* (Fig. 1D and 1E). At 3 days on SIM, continued cell division of prSPCs led to the formation of the proSAM, which comprised many cells forming a globular structure, with signals from co-expression of *STM* and *WUS* (Fig. 1D and 1E). Some psSPCs showing *WUS* expression but not *STM* expression were observed at 1 and 2 days on SIM, and those psSPCs did not further proliferate into the proSAM (Fig. 1D and 1E). From 4 to 7 days on SIM, the proSAM differentiated into the SAM, in which *WUS* expression gradually restricted into the organizing center and *STM* expression was detectable throughout the whole SAM (Fig. 1D and 1E). The SAM continuously gave rise to leaf primordia (Fig. 1D and 1E). The above data indicate that the start of *WUS* and *STM* co-expression marks the fate transition from SPCs into prSPCs (see model in Fig. 1E).

The *Arabidopsis* genome contains four class I KNOX genes, i.e. *STM, BREVIPEDICELLUS* (*BP*), *KNOTTED1-LIKE IN ARABIDOPSIS2* (*KNAT2*), and *KNAT6* (Maksimova et al., 2021). We also detected upregulation of *BP* and *KNAT2* in callus on SIM (Supplementary Table S1), and the analyses of marker lines indicated that *BP, KNAT2*, and *KNAT6* were all expressed in the prSPCs in callus culture on SIM for 2 days (Fig. 2A). To avoid the redundant functions of class I KNOX genes, we mutated the *STM* gene by Crispr/CAS9 in the *knat2-5 knat6-1 bp-9* triple mutant background to create the *bp-9 knat2-5 knat6-1 stm-ko5401* and *bp-9 knat2-5 knat6-1 stm-ko5402* quadruple mutants (Supplementary Fig. S3 and S4).

We then tested whether *STM* and class I KNOX genes are required for shoot organogenesis in tissue culture. Compared with the wild type Col-0, the *stm-6* single mutant showed partially defective shoot regeneration, and the *bp-9 knat2-5 knat6-1 stm-ko5401* and *bp-9 knat2-5 knat6-1 stm-ko5402* quadruple mutants barely produced shoots from callus on SIM (Fig. 1F and 1G). In addition, β-estradiol-induced overexpression of *STM* (*pER8:STM*) on SIM promoted shoot regeneration from callus (Fig. 1H). These findings indicate that *STM* may have partially redundant roles with class I KNOX genes in controlling shoot organogenesis in tissue culture.

Overall, the results of these experiments show that the initial expression of *STM* together with *WUS* at 1 to 2 days on SIM is the marker for the fate establishment of prSPCs from SPCs, and that *STM*, together with other class I KNOX genes, is essentially required for *de novo* shoot regeneration in *Arabidopsis* tissue culture. In addition, *STM* overexpression might serve as a candidate molecular tool that can enhance shoot regeneration in tissue culture. A previous study indicated that *CUPSHAPED COTYLEDON* (*CUC*) genes can activate *STM* expression during *de novo* shoot regeneration (Daimon et al., 2003), and *CUC2* was shown to be involved in the prSPC fate establishment (Varapparambath et al., 2022). In further research, it will be important to connect the gene network involving *CUC*s and other SAM-related genes in the fate transition from SPCs into prSPCs.

## Acknowledgments

No conflict of interest is declared. We thank Y. Jiao, V. Pautot, S. Hake, GABI-Kat, and NASC for *Arabidopsis* seeds used in this work. We thank J. Zhu and Y. Mao for providing the CRISPR/Cas9 vectors. We thank X. Fang for the assistance during the construction of *pER8:STM*. We thank Liwen Bianji for editing the English text of a draft of this manuscript.

## Author contributions

N.Z. and L.X. designed the project and wrote the manuscript; and N.Z. and D.X. performed the experiments and analyzed the data.

## Supplementary data

Supplementary Figure S1. Clusters of genes with different expression patterns.

Supplementary Figure S2. Expression patterns of *BP, KNAT2*, and *KNAT6*.

Supplementary Figure S3. T-DNA and CRISPR/Cas9 mutations in *STM*.

Supplementary Figure S4. Quadruple KNOX mutants.

Supplementary Table S1. List of gene clusters and primers.

Supplementary Materials and Methods.

## Data availability

The datasets used and/or analyzed during the current study are available from the corresponding author upon request.

## Supplementary Data

**Figure S1.**
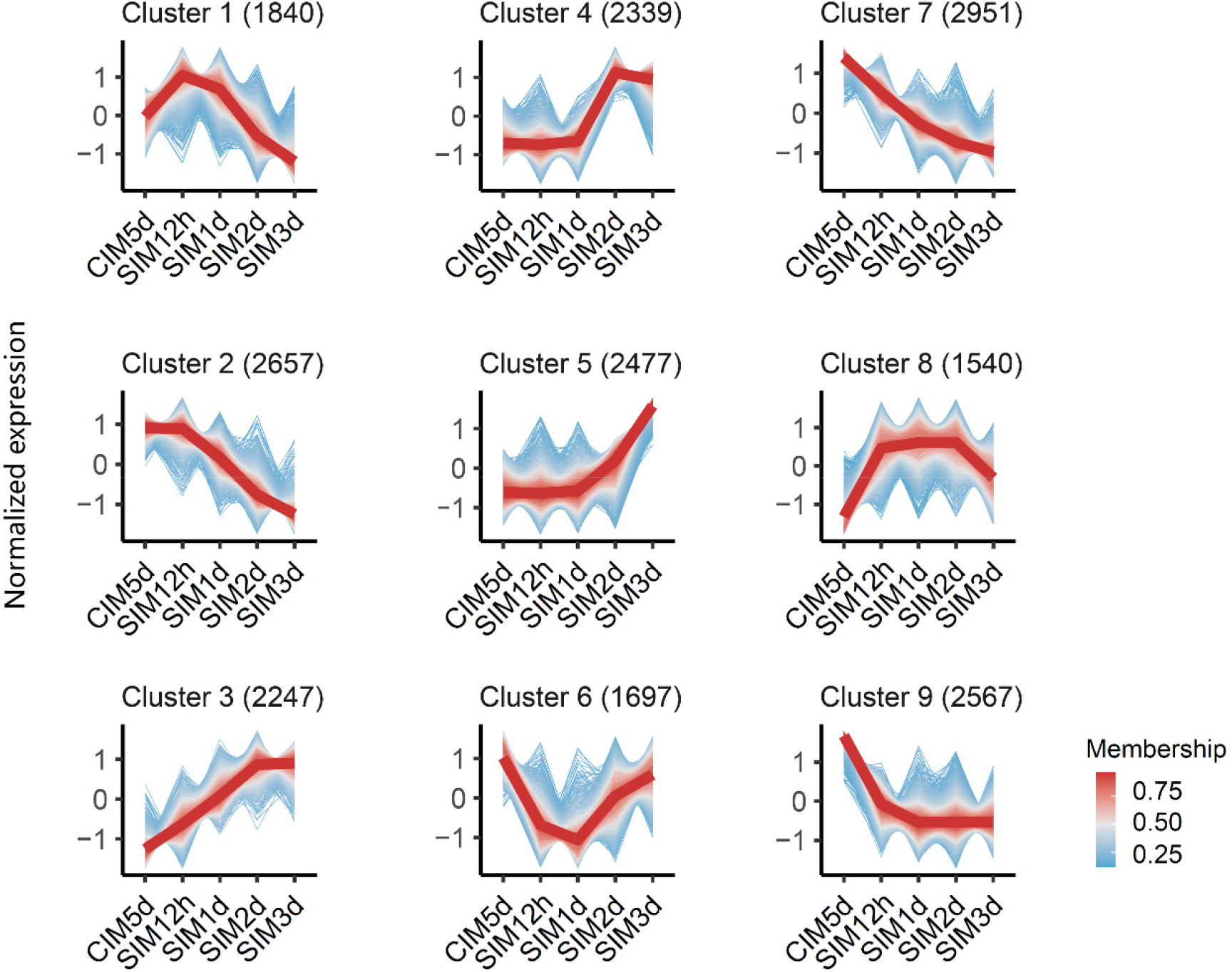
Clusters of genes with different expression patterns. Nine clusters of genes with different expression patterns in Col-0 hypocotyls during tissue culture on CIM and SIM. Clusters 8 and 6 are also shown in Figure 1B. See Table S1 for full list of genes in each cluster.

**Figure S2.**
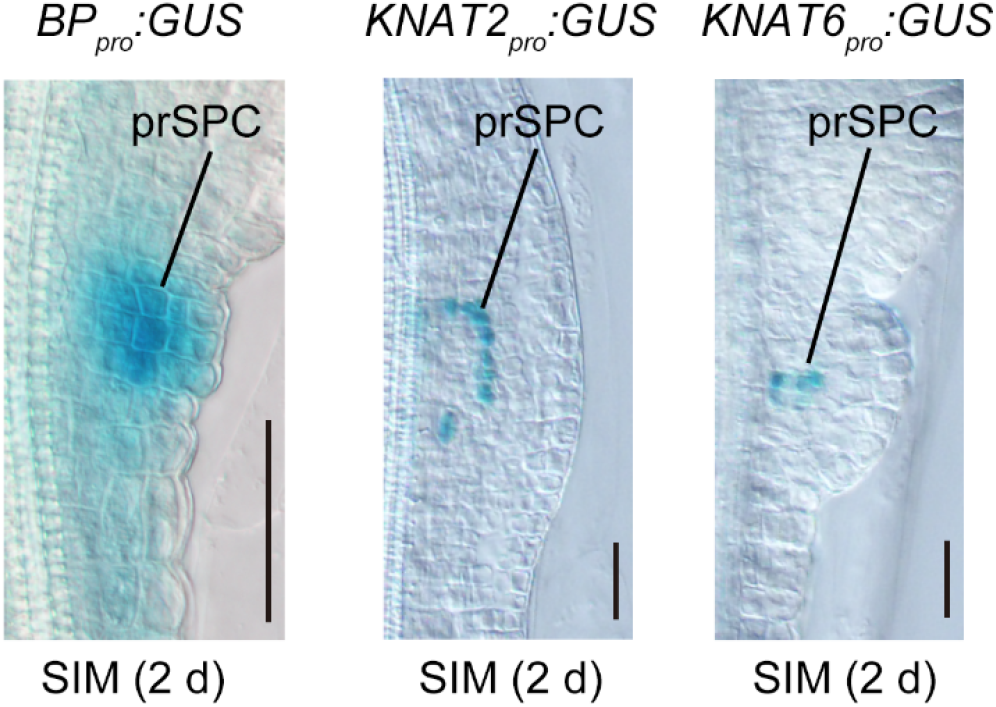
Expression patterns of *BP, KNAT2*, and *KNAT6*. GUS signals from *BP*_*pro*_*:GUS, KNAT2*_*pro*_*:GUS*, and *KNAT6*_*pro*_*:GUS* show that the three genes are expressed in the prSPCs on SIM at 2 days.

**Figure S3.**
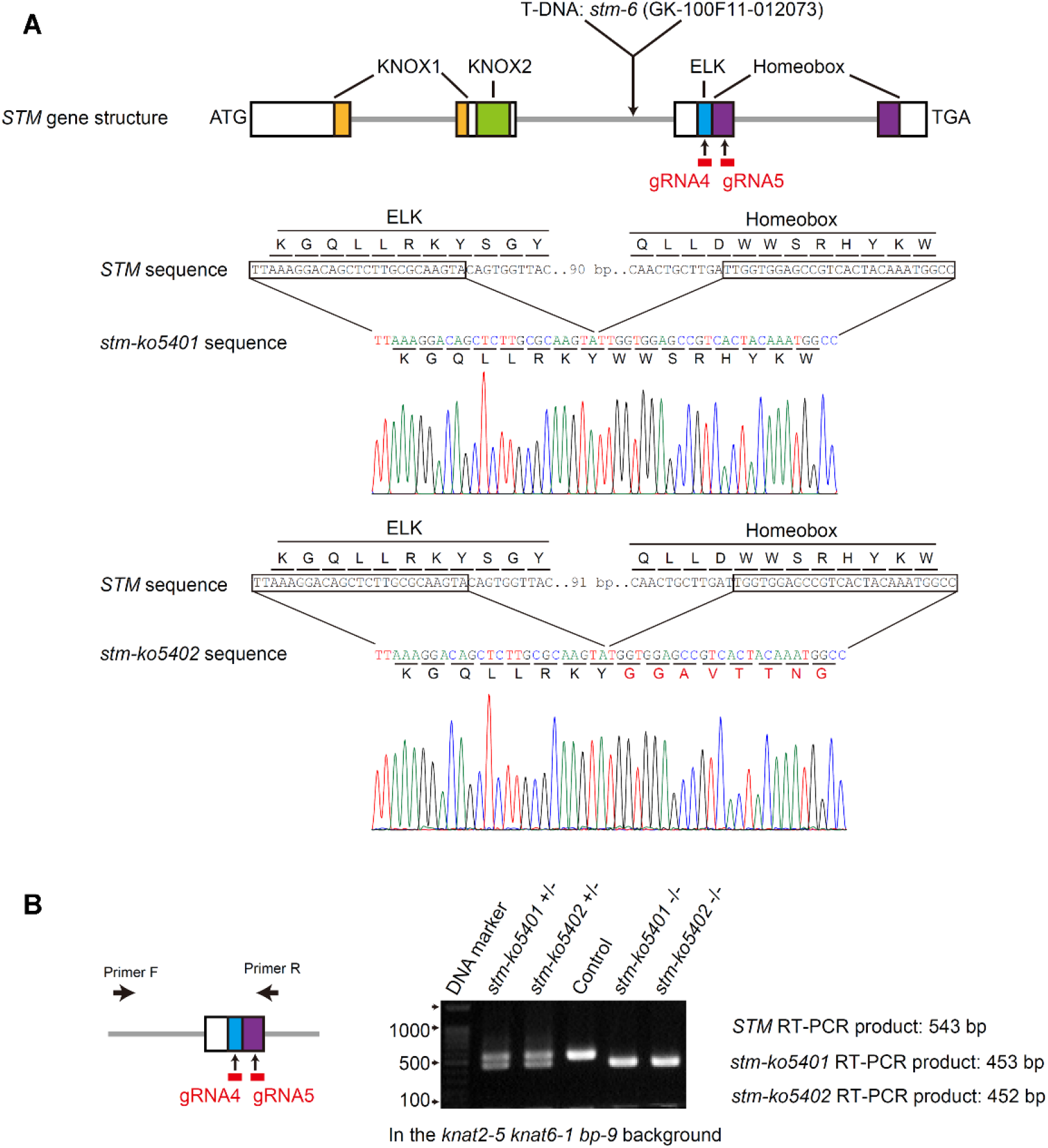
T-DNA and CRISPR/Cas9 mutations in *STM*. **(A)** Schematic of T-DNA and CRISPR/Cas9 mutations in *STM*. **(B)** Genomic PCR analysis of CRISPR/Cas9 mutations in *STM*.

**Figure S4.**
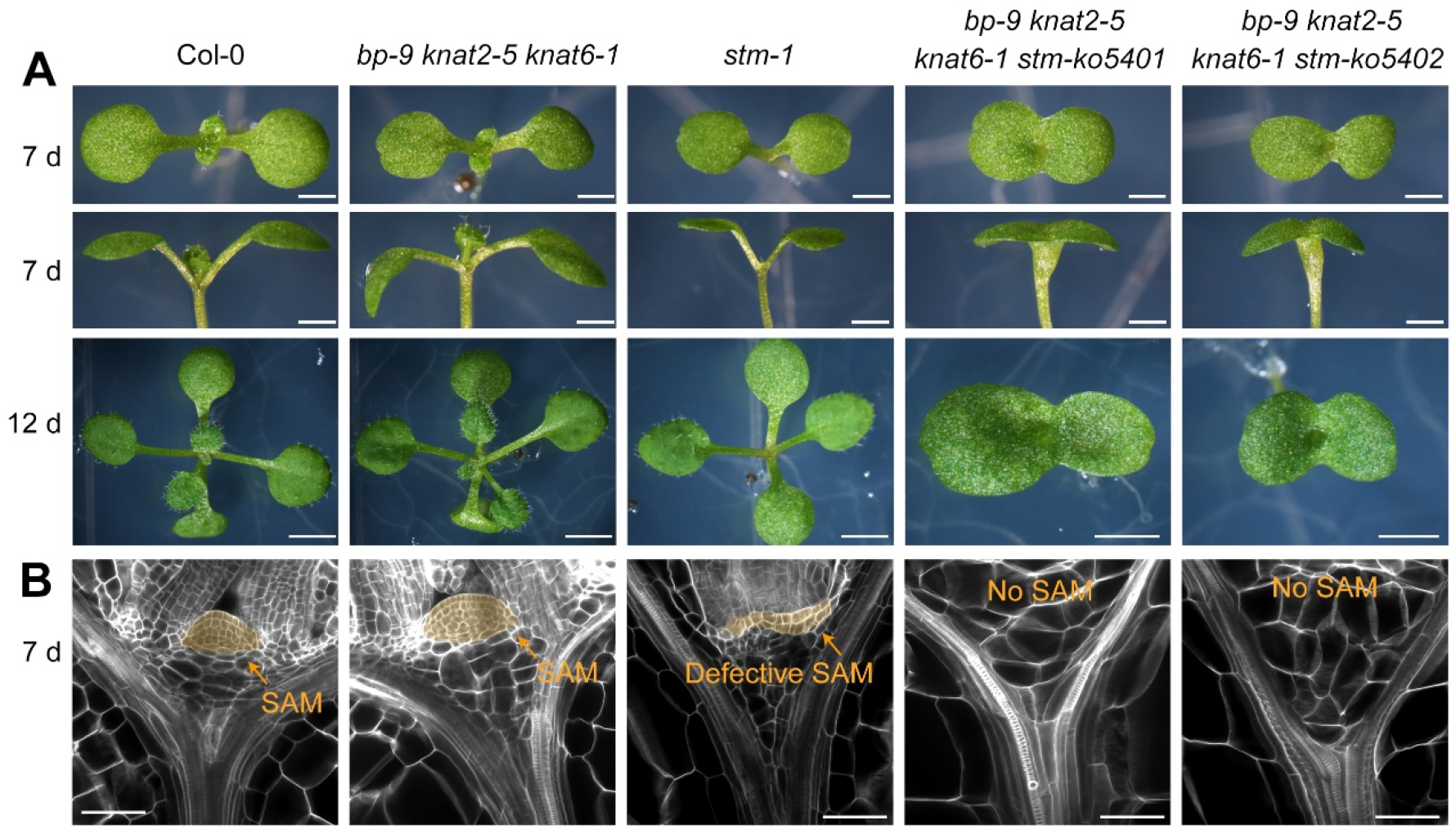
Quadruple KNOX mutants. **(A, B)** Phenotypes of seedlings (A) and ClearSee images of SAM (B) from Col-0 (wild type), *bp-9 knat2-5 knat6-1* triple mutant, *stm-6* single mutant, and *bp-9 knat2-5 knat6-1 stm-ko5401* and *bp-9 knat2-5 knat6-1 stm-ko5402* quadruple mutants. The *stm-6* single mutant showed a phenotype of defective SAM development similar to that of other *stm* mutants (Endrizzi et al., 1996; Long et al., 1996), and the *bp-9 knat2-5 knat6-1 stm-ko5401* and *bp-9 knat2-5 knat6-1 stm-ko5402* quadruple mutants showed a stronger phenotype with cup-shaped fused cotyledons and no SAM development, similar to the *knat6 stm* double mutant (Belles-Boix et al., 2006). Scale bars, 1 mm in (A) and 50 μm in (B).

**Table S1. List of gene clusters and primers.**

**(A)** List of all gene clusters in Supplemental Figure 1.

**(B)** List of primers used in this study.

## Supplementary Materials and methods

### Plant materials and culture conditions

The *bp-9 knat2-5 knat6-1* triple mutant (Li et al., 2012) and the *BP*_*pro*_*:GUS* (Ori et al., 2000), *KNAT2*_*pro*_*:GUS* (Hamant et al., 2002), and *KNAT6*_*pro*_*:GUS* (Belles-Boix et al., 2006) lines have been described previously. *stm-6* is a GABI-Kat T-DNA insertion line (GK-100F11-012073) (Kleinboelting et al., 2012) (Supplementary Figure 2). *STM*_*pro*_*:gSTM-Venus* was described previously (Cao et al., 2020).

The *WUS*_*pro*_*:2×mCherry-ER* vector was constructed by inserting the 5.1-kb upstream promoter of *WUS, 2×mCherry-ER*, and the 3.1-kb downstream sequence of *WUS* into the *pBI101* vector. The *pER8:STM* was constructed by inserting the *STM* cDNA into the *pER8* vector (Zuo et al., 2000). These constructs were introduced into Col-0 by *Agrobacterium tumefaciens*-mediated transformation.

To generate the *stm-ko5401* and *stm-ko5402* mutants (Supplementary Figure 2), *STM*-specific sequences (5′ - GTCAACAACTGCTTGATTGG **TGG** (PAM sequence) - 3′ and 5′ - CAGCTCTTGCGCAAGTACAG **TGG** (PAM sequence) - 3′) were selected as the target for Cas9 to mutate *STM*. Vectors for CRISPR/Cas9 were constructed as previously described (Liu et al., 2015). The constructs were introduced into the *bp-9 knat2-5 knat6-1* triple mutant background by *A. tumefaciens*-mediated transformation. For *de novo* shoot regeneration in tissue culture, *Arabidopsis* seeds were grown vertically on ½ Murashige and Skoog (MS) medium (1.2% w/v agar) in the light for 1 day and then in the dark for 8 days to allow hypocotyls to elongate. Approximately 1-cm hypocotyl segments were cut and cultured on CIM (MS medium, 2% w/v sucrose, pH 5.8, 0.8% w/v agar, 11 μM 2,4-dichlorophenoxyacetic acid, and 0.2 μM kinetin) under continuous light conditions for 5 days to induce callus formation. The calli were then transferred to SIM [MS medium, 2% w/v sucrose, pH 5.7, 0.8% w/v agar, 2 μM 6-(dimethylallylamino) purine, 0.9 μM indole acetic acid] and cultured under continuous light conditions for 7 days to induce shoot organogenesis.

The primers used in this study are listed in Supplementary Table 1.

### ClearSee assay and confocal microscopy

For observation of fluorescent marker lines, the ClearSee assay was performed as previously described (Kurihara et al., 2015; Zhai and Xu, 2021). The various tissues were observed and photographed using an Eclipse Ti confocal microscope (Nikon).

### RNA-seq analysis

For RNA-seq analysis, total RNA was extracted from calli collected from CIM and SIM at various times (CIM 5 d, SIM 12 h, SIM 24 h, and SIM 48 h) using the TRIzol method. Sequencing was performed by LC Sciences (Hangzhou, China) on the Illumina Hiseq platform (Illumina, San Diego, CA, USA). Raw data were processed to obtain TPM data (Dobin et al., 2013). Clustering of gene expression patterns across different sampling time points was conducted using the mfuzz method in ClusterGVis (https://github.com/junjunlab/ClusterGVis).

The raw RNA-seq data have been deposited in the Genome Sequence Archive (Genomics, Proteomics & Bioinformatics 2021) at the National Genomics Data Center, China National Center for Bioinformation/Beijing Institute of Genomics, Chinese Academy of Sciences (https://ngdc.cncb.ac.cn/gsa) under the accession number GSA: CRA023698. The RNA-seq data can also be accessed using the online tool (http://xulinlab.cemps.ac.cn/).

### Gene sequence data

Sequence data relevant to this article can be found in the *Arabidopsis* Genome Initiative under the following accession numbers: *WUS* (AT2G17950), *STM* (AT1G62360), *BP* (AT4G08150), *KNAT2* (AT1G70510), and *KNAT6* (AT1G23380).

